# RAG1 and RAG2 Non-core Regions Are Implicated in Leukemogenesis and Off-target V(D)J Recombination in BCR-ABL1-driven B-cell Lineage Lym-phoblastic Leukemia

**DOI:** 10.1101/2023.09.29.560132

**Authors:** Xiaozhuo Yu, Wen Zhou, Xiaodong Chen, Shunyu He, Mengting Qin, Meng Yuan, Yang Wang, Woodvine otieno Odhiambo, Yinsha Miao, Yanhong Ji

## Abstract

The evolutionary conservation of non-core RAG regions suggests significant roles that might involve quantitative or qualitative alterations in RAG activity. Off-target V(D)J recombination contributes to lymphomagenesis and is exacerbated by RAG2’ C-terminus absence in Tp53^-/-^ mice thymic lymphomas. However, the genomic stability effects of non-core regions from both cRAG1 and cRAG2 in *BCR-ABL1*^+^ Blymphoblastic leukemia (*BCR-ABL1*^+^ B-ALL), the characteristics, and mechanisms of non-core regions in suppressing off-target V(D)J recombination remains unclear. Here, we established three mouse models of *BCR-ABL1*^+^ B-ALL in mice expressing full-length RAG (fRAG), core RAG1 (cRAG1), and core RAG2 (cRAG2). The cRAG (cRAG1 and cRAG2) leukemia cells exhibited greater malignant tumor characteristics compared to fRAG cells. Additionally, cRAG cells showed higher frequency of off-target V(D)J recombination and oncogenic mutations than fRAG. We also revealed decreased RAG cleavage accuracy in cRAG cells and a smaller recombinant size in cRAG1 cells, which could potentially exacerbate off-target V(D)J recombination in cRAG cells. In conclusion, these findings indicate that the non-core RAG regions, particularly the non-core region of RAG1, play a significant role in preserving V(D)J recombination precision and genomic stability in *BCR-ABL1*^+^ B-ALL.

## Introduction

V(D)J recombination serves as the central process for early lymphocyte development and generates diversity in antigen receptors. This process involves the double-strand DNA breaks of gene segments by the V(D)J recombinase, including RAG1 and RAG2. RAG recognizes conserved recombination signal sequences (RSSs) positioned adjacent to V, D, and J gene segments. A bona fide RSS contains a conserved palindromic heptamer (consensus 5′-CACAGTG) and A-rich nonamer (consensus 5′-ACAAAAACC) separated by a degenerate spacer of either 12 or 23 base pairs (***Hirokawa, et al., 2020; Schatz and Ji, 2011)***. The process of efficient recombination is contingent upon the presence of recombination signal sequences (RSSs) with differing spacer lengths, as dictated by the “12/23 rule” (***Banerjee and Schatz, 2014;Eastman, et al., 1996;Shi, et al., 2020)***. Following cleavage, the DNA ends are joined via non-homologous end joining (NHEJ), resulting in the precise alignment of the two coding ends and the signal ends (***Rooney, et al., 2004)***. V(D)J recombination promotes B cell development, but aberrant V(D)J recombination can lead to precursor B-cell malignancies through RAG mediated off-target effects (***Mendes, et al., 2014;Onozawa and Aplan, 2012;Thomson, et al., 2020)***.

The regulation of RAG expression and activity is multifactorial, serving to ensure V(D)J recombination and B cell development (***Gan, et al., 2021;Kumari, et al., 2021)***. The RAGs consist of core and non-core region. Although non-core regions of RAG1/2 are not strictly required for V(D)J recombination, the evolutionarily conserved non-core RAG regions indicate their potential significance in vivo that may involve quantitative or qualitative modifications in RAG activity and expression (***Braams, et al., 2023;Curry and Schlissel, 2008;Liu, et al., 2022;Liu, et al., 2022;Sekiguchi, et al., 2001)***. Specifically, the non-core RAG2 region (amino acids 384– 527 of 527 residues) contains a plant homeodomain (PHD) that can recognize histone H3K4 trimethylation, as well as a T490 locus that mediates a cell cycle-regulated protein degradation signal in proliferated pre-B cells stage (***Liu, et al., 2007;***

***Matthews, et al., 2007)***. Failure to degrade RAG2 during the S stage poses a threat to the genome (***Zhang, et al., 2011)***. Moreover, the off-target V(D)J recombination frequency is significantly higher when RAG2 is C-terminally truncated, thereby establishing a mechanistic connection between the PHD domain, H3K4me3-modified chromatin, and the suppression of off-target V(D)J recombination (***Lu, et al., 2015;Mijuškovi***ć***, et al., 2015)***. The RAG1’ non-core region (amino acids1–383 of 1040 residues) has been identified as a RAG1 regulator. While the core RAG1 maintains its catalytic activity, it’s in vivo recombination efficiency and fidelity are reduced in comparison to the full-length RAG1 (fRAG1). In addition, the RAG1 binding to the genome is more indiscriminate (***Beilinson, et al., 2021;Sadofsky, et al., 1993)***. The N-terminal domain (NTD), which is evolutionarily conserved, is predicted to contain multiple zinc-binding motifs, including a Really Interesting New Gene (RING) domain (aa 287 to 351) that can ubiquitylate various targets, including RAG1 itself (***Deng, et al., 2015)***.. Furthermore, NTD contains a specific region (amino acids 1 to 215) that facilitates interaction with DCAF1, leading to the degradation of RAG1 in a CRL4-dependent manner (***Schabla, et al., 2018)***. Additionally, the NTD plays a role in chromatin binding and the genomic targeting of the RAG complex (***Schatz and Swanson, 2011)***. Despite increased evidence emphasizing the significance of non-core RAG regions, particularly RAG1’s non-core region, the function of non-core RAG regions in off-target V(D)J recombination and the underlying mechanistic basis have not been fully clarified.

Typically, genomic DNA is safeguarded against inappropriate RAG cleavage by the inaccessibility of cryptic RSSs (cRSSs), which are estimated to occur once per 600 base pairs (***Teng, et al., 2015)***. However, recent research has demonstrated that epigenetic reprogramming in cancer can result in heritable alterations in gene expression, including the accessibility of cRSSs (***Becker, et al., 2020;Fatma, et al., 2022;Goel, et al., 2022;Khoshchehreh, et al., 2019)***. We selected the *BCR-ABL1*^+^ B-ALL model, which is characterized by ongoing V(D)J recombinase activity and *BCR-ABL1* gene rearrangement in pre-B leukemic cells (***Schjerven, et al., 2017;Wong and Witte, 2004)***. The genome structural variations (SVs) analysis was conducted on leukemic cells from *fRAG, cRAG1*, and *cRAG2, BCR-ABL1*^+^ B-ALL mice to examine the involvement of non-core RAG regions in off-target V(D)J recombination events. The non-core domain deletion in both *RAG1* and *RAG2* led to accelerated leukemia onset and progression, as well as an increased off-target V(D)J recombination. Our analysis showed a reduction in RAG cleavage accuracy in *cRAG* cells and a decrease in recombinant size in *cRAG1* cells, which may be responsible for the increased off-target V(D)J recombination in *cRAG* leukemia cells. In conclusion, our results highlight the potential importance of the non-core RAG region, particularly RAG1’s non-core region, in maintaining accuracy of V(D)J recombination and genomic stability in *BCR-ABL1*^+^ B-ALL.

## Results

### cRAG give more aggressive leukemia in a mouse model of *BCR-ABL1*^+^ B-ALL

In order to assess the impact of RAG activity on the clonal evolution of *BCR-ABL1^+^* B-ALL through a genetic experiment, we utilized bone marrow transplantation (BMT) to compare disease progression in fRAG, cRAG1, and cRAG2 *BCR-ABL1^+^* B-ALL (***Schjerven, et al., 2017;Wong and Witte, 2004)***. Bone marrow cells transduced with a BCR-ABL1/GFP retrovirus were administered into syngeneic lethally irradiated mice, and CD19^+^ B cell leukemia developed within 30-80 days (Figure 1A, Supplementary Figure 1). Western blotting results confirmed equivalent transduction efficiencies of the retroviral *BCR-ABL1* in all three cohorts (Supplementary Figure 2A). To investigate potential variances in leukemia outcome across different genomic backgrounds, we employed Mantel-Cox analysis to evaluate survival rates in fRAG, cRAG1, or cRAG2 mice transplanted with *BCR-ABL1*-transformed bone marrow cells. Our results show that, during the primary transplant phase, *BCR-ABL1*^+^ B-ALL mice expressing cRAG1 or cRAG2 demonstrated lower survival rates compared to their counterparts with fRAG (median 74.5 days versus 39 or 57 days, *P* < 0.0425, Figure 1A). This survival rates discrepancy was also observed during the secondary transplant phase, wherein leukemic cells were extracted from the spleens of primary recipients and subsequently purified via GFP^+^ cell sorting. A total of 10^5^,10^4^ and 10^3^ GFP^+^ leukemic cells that originated from fRAG, cRAG1, or cRAG2 leukemic mice were transplanted into corresponding non-irradiated immunocompetent syngenetic recipients (survival days fRAG,11-26 days, cRAG1,10-16 days, cRAG2,11-21 days, Supplementary Figure 2B). Additionally, the cRAG mice exhibited significantly higher leukemia burdens in the peripheral blood, bone marrow, and spleen compared to the fRAG mice (Figure 1B-D). To elucidate the cellular mechanisms driving the accelerated proliferation observed in cRAG *BCR-ABL1*^+^ B-ALL, flow cytometry analyses were conducted to evaluate cell cycle dynamics and apoptotic activity. Results revealed a higher fraction of cRAG *BCR-ABL1*^+^ B-ALL cells residing in the S/G2-M phase of the cell cycle compared to their fRAG counterparts (Figure 1E). Additionally, the enhanced proliferation in cRAG leukemic cells was attributed to a reduction in apoptosis rates (Supplementary Figure 2C). RNA-seq analysis demonstrated the changes of cell differentiation and proliferation/apoptotic pathways (Supplementary Figure 3) These findings indicate that the absence of non-core RAG regions accelerates malignant transformation and leukemic proliferation, leading to a more aggressive disease phenotype in the cRAG *BCR-ABL1*^+^ B-ALL mouse model.

**Figure 1:**
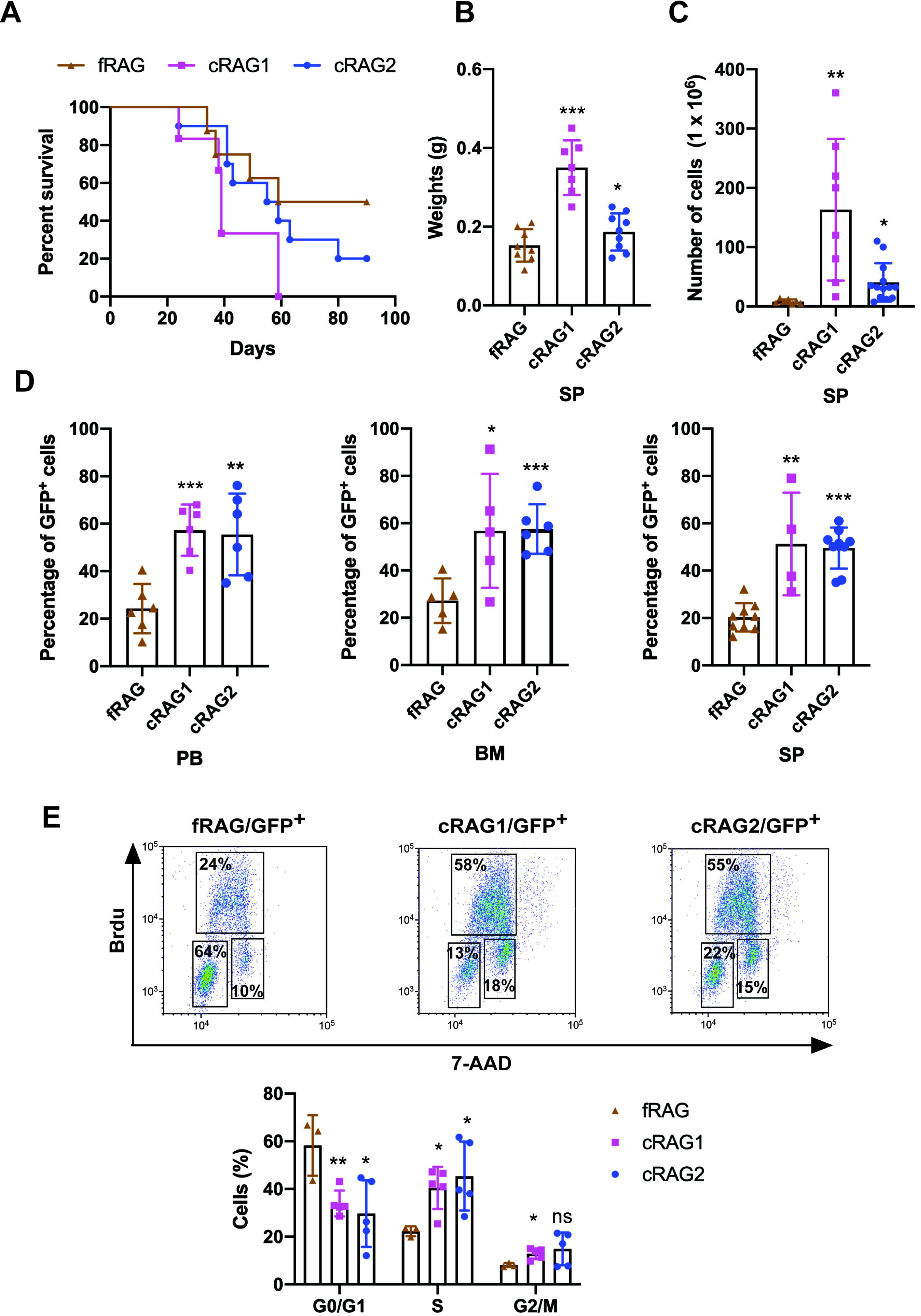

**Figure 2:**
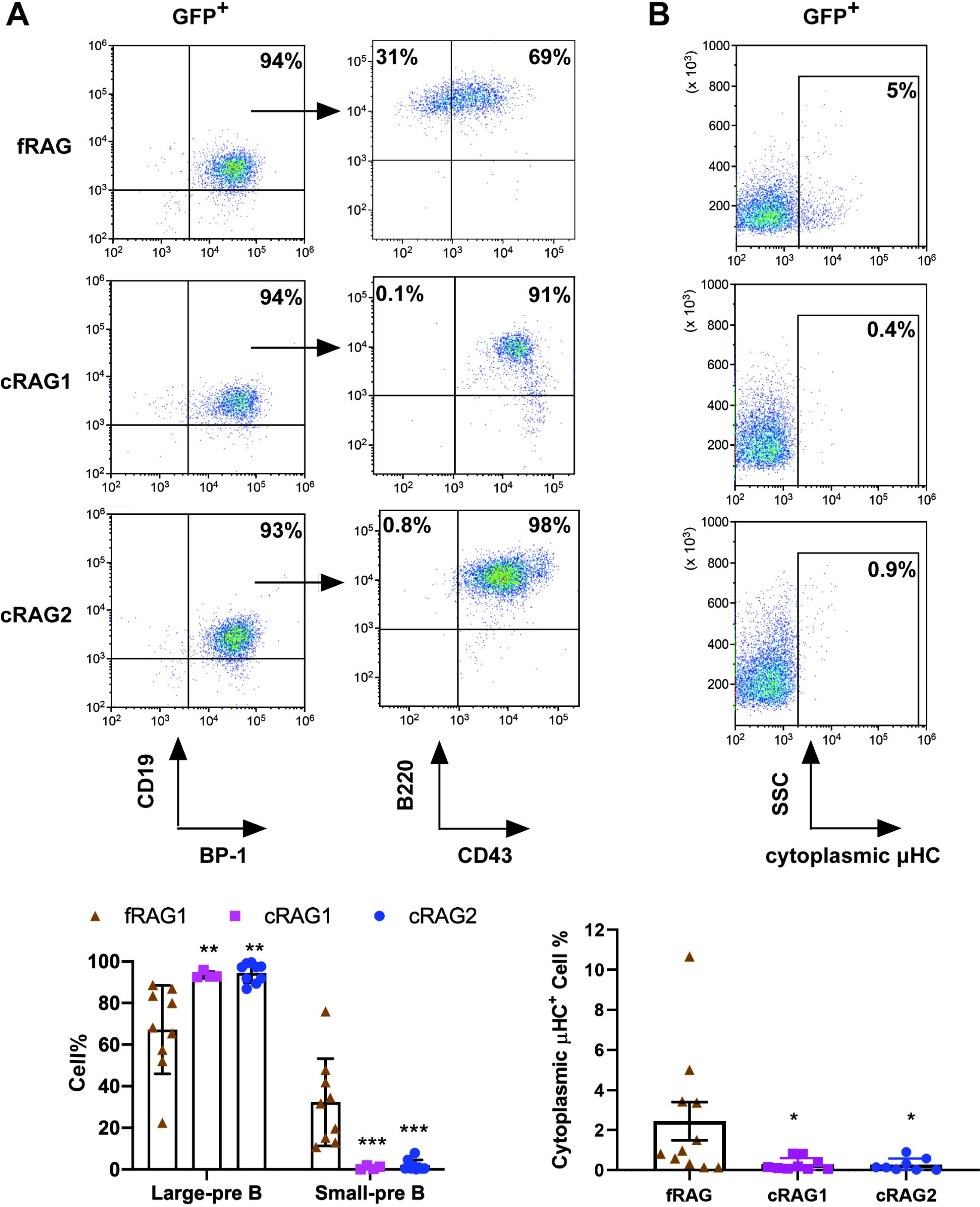

### The loss of non-core RAG regions corresponds to a less mature cell surface phenotype but does not impede IgH VDJ recombination

To delineate the developmental stages of B cells from which the leukemic cells originated, we performed flow cytometry on single cells stained with B cell-specific surface markers. Analysis revealed that 91%-98% of GFP^+^ cells in cRAG mice were CD19^+^BP-1^+^B220^+^CD43^+^, indicating that most leukemic cells were at the large pre-B cell stage (Figure 2A) (***Hardy and Hayakawa, 2001)***. Conversely, in fRAG leukemic mice, the distribution was 65% large pre-B cells (GFP^+^CD19^+^BP-1^+^B220^+^CD43^+^) and 35% small pre-B cells (GFP^+^CD19^+^BP-1^+^B220^+^CD43^-^) (Figure 2A). Moreover, approximately 5% of leukemic cells in fRAG mice expressed μHC, in contrast to minimal expression in cRAG leukemic cells. This suggests that fRAG leukemic cells may differentiate further, associated with an immune phenotype (Figure 2B). *IgH* rearrangement initiates with D_H_-J_H_ joining in pro-B cells, followed by V_H_-DJ_H_ joining in pro-B cells, and ultimately, V_L_-J_L_ rearrangements occur at the *IgL* loci in small pre-B cells (***Schatz and Ji, 2011)***. Genomic PCR analysis of DNA from GFP^+^CD19^+^ cells was utilized to assess V_H_DJ_H_ rearrangement. The results showed a pronounced oligoclonality in cRAG leukemic cells, with tumors consistently demonstrating rearrangements involving a restricted set of V_H_ family members. In contrast, fRAG leukemias displayed significant polyclonality, evidenced by the widespread rearrangement of various V_H_ family members to all potential J_H_1-3 segments, indicative of a broader clonal diversity (Supplementary Figure 4). This observation aligns with the more aggressive leukemia phenotype seen in cRAG *BCR-ABL1*^+^ B-ALL mice. Such oligoclonality in cRAG leukemic cells suggests a selection process driven by BCR-ABL1-induced leukemia, favoring the emergence of a limited number of dominant leukemic clones. The absence of non-core RAG regions appears to restrict the diversity of leukemic clones, leading to the formation of oligoclonal tumors.

### The loss of non-core RAG regions highlights genomic DNA damage

The findings indicate that leukemic cells from three types of mice exhibited variable arrests at the large pre-B cell stage, deviating from normal B cell developmental trajectory. Typically, at this juncture, B cells initiate the degradation of RAG2 via the cyclin-dependent kinase cyclinA/Cdk2, leading to a downregulation of RAG activity. It is therefore crucial to explore the impact of deletions in non-core regions on the expression and functionality of RAG in these leukemic cells. Analysis showed that both RAG1 (cRAG1) and RAG2 (cRAG2) were present in GFP^+^CD19^+^ splenic leukemic cells from *BCR-ABL1*^+^ B-ALL mice across different genetic backgrounds (Figure 3A). Notably, we observed an upregulation of cRAG1 and cRAG2 in leukemic cells from cRAG1 or cRAG2 mice compared to those from fRAG mice (Figure 3A, Supplementary Figure 5A). The in vitro V(D)J recombination assay confirmed that different forms of RAG exhibited cleavage activity in *BCR-ABL1*^+^B-ALL (Figure 3B and Supplementary Figure 5B).

**Figure 3:**
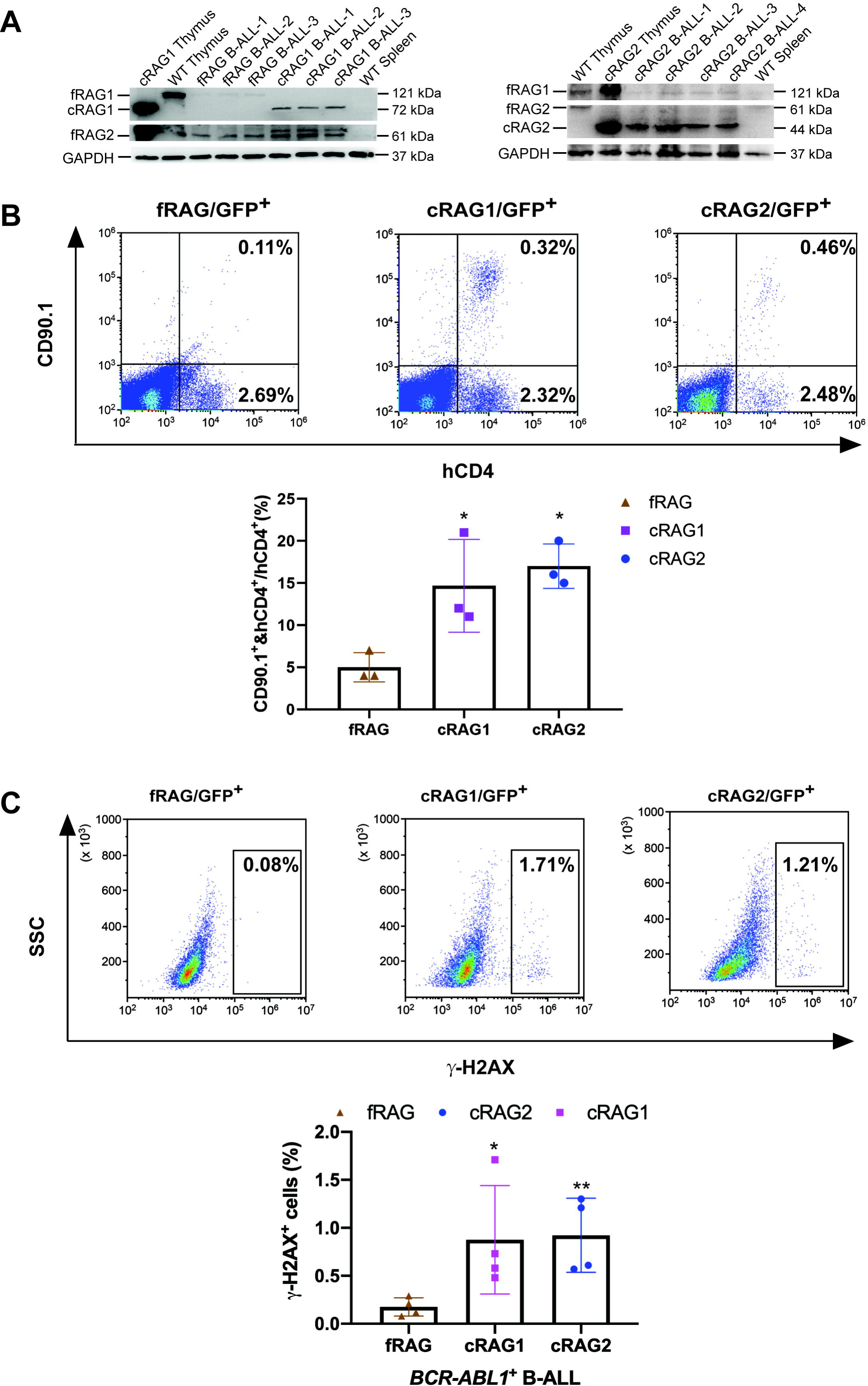

To examine the potential correlation between aberrant RAG activity and increased DNA double-strand breaks (DSBs), we assessed levels of phosphorylated H2AX (L-H2AX), a marker of DSB response, in leukemic cells from fRAG, cRAG1, and cRAG2 mice (gated on GFP^+^). This evaluation aimed to gauge DNA DSBs and overall genomic instability. Flow cytometry analysis revealed elevated L-H2AX levels in cRAG1 and cRAG2 leukemic cells compared to those from fRAG, indicating a more pronounced role of cRAG in mediating somatic structural variants in *BCR-ABL1*^+^ B cells. These findings suggest enhanced expression of cRAG1 endonucleases in cRAG1 leukemic cells and increased DNA damage in cells lacking core RAG regions.

### Off-target recombination mediated by RAG in *BCR-ABL1*^+^ B cells

Genome-wide sequencing and analysis were performed to compare somatic structural variants (SVs) in *BCR-ABL1*^+^ B cells derived from fRAG, cRAG1 and cRAG2 mice. The leukemic cells were sequenced with an average coverage of 25× (Supplementary Table 1). The SVs generated by RAG were screened based on two criteria: the presence of a CAC to the right (or GTG to the left) of both breakpoints, and its occurrence within 21 bp from the breakpoint (***Mijuškovi***ć***, et al., 2015)***. Further elaboration on these criteria can be found in Supplementary Figure 6. Consequently, aberrant *V*-to-*V* junctions and *V* to intergenic regions were encompassed in five validated abnormal rearrangements at *Ig* loci in cRAG leukemic mice (Supplementary Table 2). Additionally, seven samples had 24 somatic structural variations, with an average of 3.4 coding region mutations per sample (range of 0-9), which is consistent with the limited number of acquired somatic mutations observed in hematological cancers (Figure 4 and Supplementary Table 3). The results of the study demonstrate that fRAG cells had low SVs (0-1 per sample), cRAG1 cells exhibited higher SVs (6-9 per sample) while cRAG2 cells had moderate SVs incidence (1-4 per sample) (Figure 4, Supplementary Table 3). These findings suggest that cRAG may lead to an elevated off-target recombination, eventually posing a threat to the *BCR-ABL1*^+^ B cells genome.

**Figure 4:**
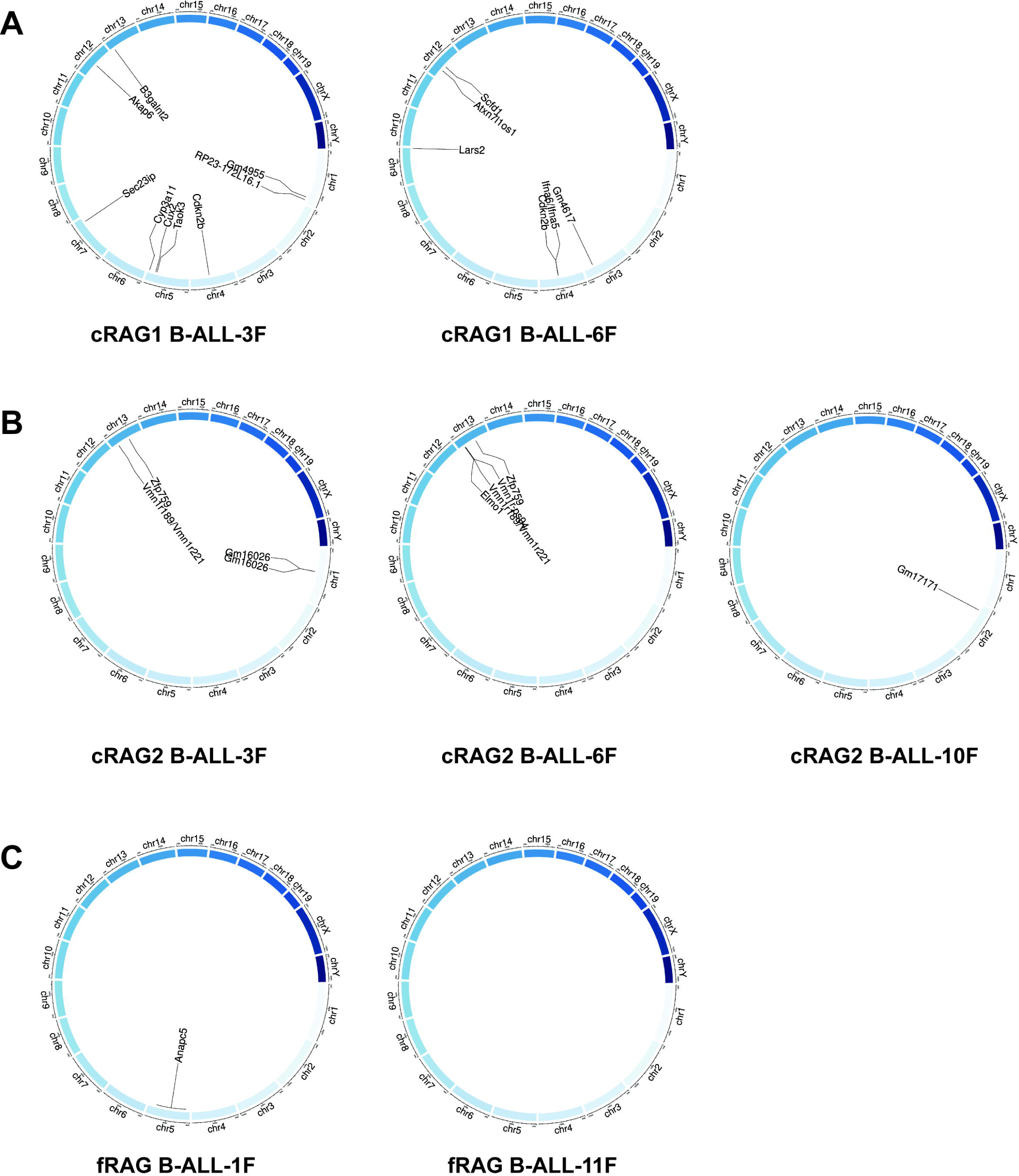

**Figure 5:**
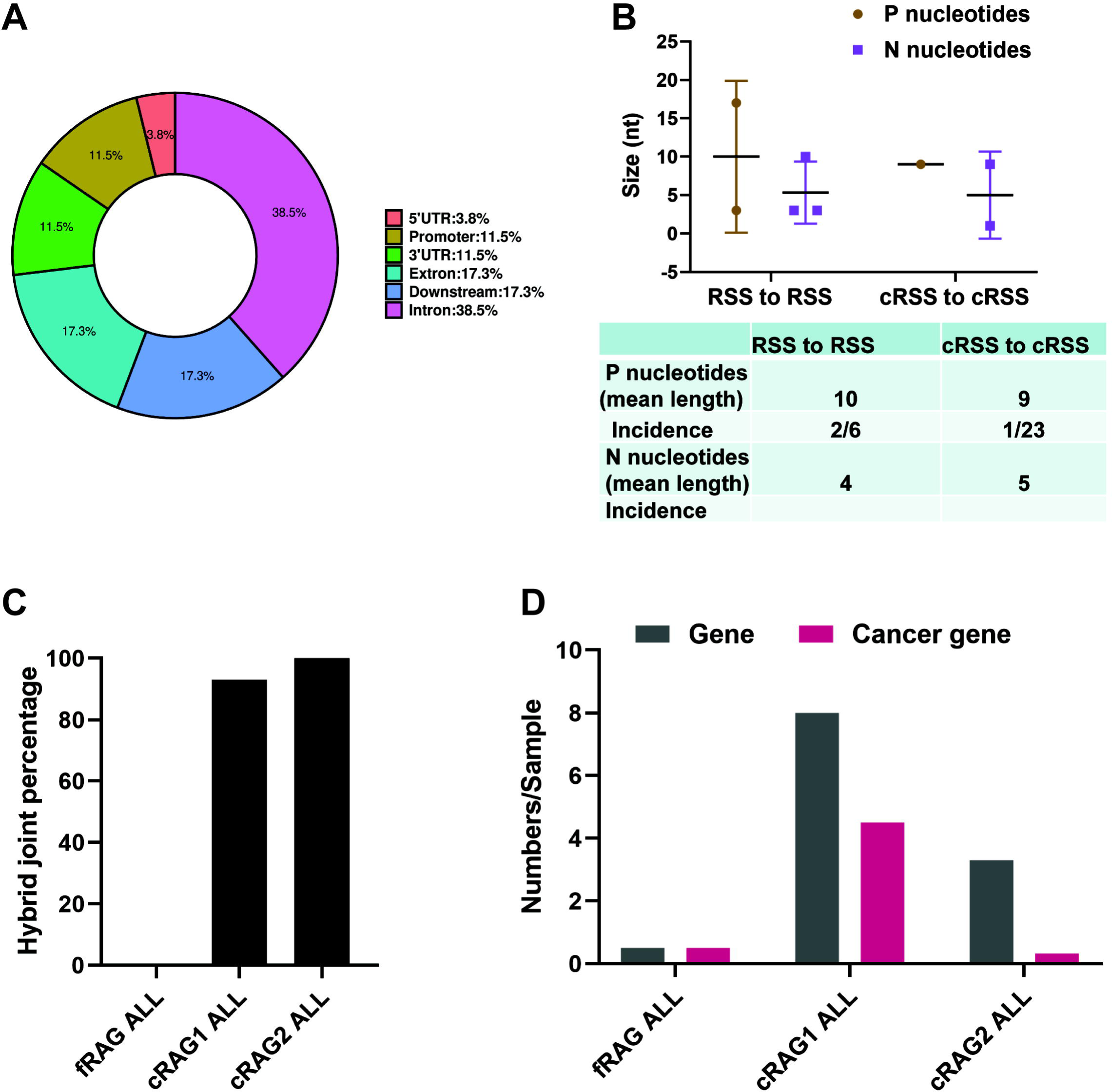

### Off-target V(D)J recombination characteristics in *BCR-ABL1^+^*B cells

We further examined the characteristics of the identified structural variants (SVs). Specifically, we analyzed the exon-intron distribution profiles of 41 breakpoints from 24 SVs through genome analysis. The results indicated that 57% of the breakpoints were located within the gene body, while 43% were enriched in the flanking sequences, the majority of which were identified as transcriptional regulatory sequences (Figure 5A). P and N nucleotides are recognized as distinctive characteristics of V(D)J recombination (***Repasky, et al., 2004)***. RSS-to-RSS and cRSS-to-cRSS recombination have P nucleotides lengths of 7 and 9, respectively, and N lengths of 5, so nucleotide lengths are basically the same during RSS-to-RSS and cRSS-to-cRSS recombination (Figure 5B). However, the frequency of P and N sequences in RSS-to-RSS recombination was 50%/50% (P/N), compared to 4%/8% (P/N) in cRSS-to-cRSS recombination (Figure 5B). This significant reduction in the frequency of P and N sequences suggests that DNA repair at off-target sites in *BCR-ABL1*^+^ B cells diverges from the classical V(D)J re-combination repair process.

The hybrid joints were notably prevalent in cRAG1 and cRAG2 leukemic cells (93% and 100%, respectively), suggesting that the non-core regions of RAG may play a role in inhibiting harmful transposition events (Figure 5C). To evaluate the effect of deleting non-core RAG regions on the emergence of oncogenic mutations, we performed a comparative analysis of cancer-related genes across three types of leukemic cells. We found that cRAG1 leukemic cells harbored a significantly higher number of cancer genes compared to the other groups. This finding corresponds with the most aggressive leukemia phenotype observed in cRAG1 *BCR-ABL1*^+^ B-ALL mice and associated changes in their transcription profiles.

### The non-core regions have effects on RAG cleavage and off-target recombination size in *BCR-ABL1^+^* B cells

Sequence logos were employed to visually contrast RSS and cRSSs within *Ig* and non-*Ig* loci, respectively. Notably, the RSS elements in *Ig* loci displayed a higher similarity to the canonical RSS, especially at critical functional sites. The initial four nucleotides (highlighted) of the canonical heptamer sequence <u>CACA</u>GTG were recognized as the cleavage site for fRAG. Conversely, in leukemic cells, the cleavage site for cRAG was pinpointed to the first three nucleotides, the CAC trinucleotide, of the heptamer sequence (Figure 6A). While both motifs (CACA and CAC) align with the highly conserved segment of the RSS heptamer sequence, differences in the cRSS sequences across off-target genes in both fRAG and cRAG mice suggest that deletion of RAG’s non-core regions broadens the spectrum of off-target substrates in *BCR-ABL1*^+^ B cells.

**Figure 6:**
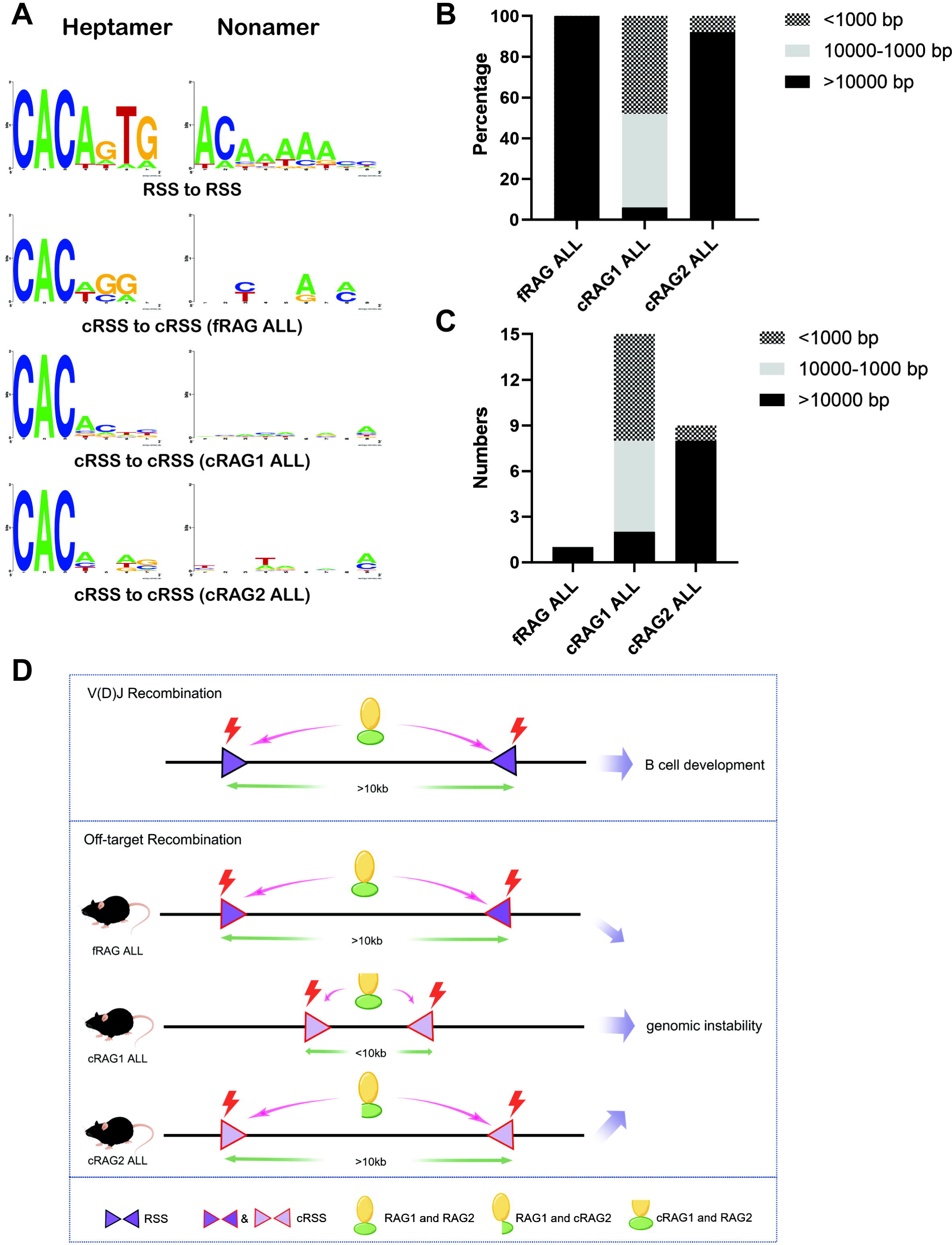

Antigen receptor genes are assembled by large-scale deletions and inversions (***Teng, et al., 2015)***. The off-target recombination size was determined as the DNA fragment size spanning the two breakpoints. Our analysis demonstrated that both fRAG and cRAG2 leukemic cells produced off-target recombinations with 100% and 92% of events, respectively, spanning over 10,000 bp in length. In contrast, cRAG1 leukemic cells showed only 6% of off-target recombinations exceeding 10,000 bp, with 48% under 1,000 bp and 46% ranging between 1,000 to 10,000 bp (Figure 6BC). These findings suggest that the cRAG1 variant primarily facilitates smaller-scale off-target recombinations in *BCR-ABL1*^+^ B cells, highlighting the role of the non-core RAG1 region in influencing the extent of off-target recombination. The deletion of the non-core RAG1 region appears to constrict the size of off-target recombination, potentially contributing to the elevated frequency of off-target V(D)J recombination observed in cRAG1 leukemic cells (Figure 6D).

## Discussion

In this study, we have demonstrated that non-core region deletion of both RAG1 and RAG2 leads to accelerated development of leukemia and increased off-target V(D)J recombination in mice models of *BCR-ABL1*^+^ B cells. Furthermore, we report reduced cRAG cleavage accuracy and off-target recombination size in cRAG leukemia cells, which might contribute to exacerbated off-target V(D)J recombination of cRAG *BCR-ABL1*^+^ B cells. These findings suggest that the non-core regions, particularly the non-core region of RAG1, play a crucial role in maintaining accuracy of V(D)J recombination and genomic stability in *BCR-ABL1*^+^ B cells.

Our findings suggest that leukemic cells with cRAG regions exhibit increased production of hybrid joints, implying that non-core RAG regions might suppress the formation of these hybrid joints in vivo. Post-cleavage synaptic complexes (PSCs), comprising RAG proteins, coding ends, and RSS ends, are believed to have evolved to form with optimal conformation and/or stability for conventional coding and RSS end-joining(***Fugmann, et al., 2000;Libri, et al., 2021)***. In contrast, cRAG PSCs could promote RAG-mediated hybrid joints by facilitating closer proximity of coding and RSS ends or by increasing PSC stability. It is also conceivable that fRAGs recruit disassembly/remodeling factors to PSCs, a process that could allow non-homologous end joining (NHEJ) factors to complete the normal reaction(***Fugmann, et al., 2000)***.

Conversely, cRAGs may have a diminished recruitment capacity due to changes in overall conformation or the absence of specific motifs, leading to more unstable PSCs and a heightened risk of accumulating incomplete hybrid joints (***Raghavan, et al., 2006;Talukder, et al., 2004)***. Our data reveal that over 90% of junctions were hybrid joints in cRAG leukemic cells, a frequency exceeding that reported in previous studies. This suggests that deficiencies in the NHEJ pathway could contribute to chromosomal instability and lymphomagenesis(***Gaymes, et al., 2002;Rassool, 2003;Scully, et al., 2019;Wiegmans, et al., 2021)***. Significantly, our analysis uncovered variations in the NHEJ repair pathway among leukemic cells from different genetic backgrounds, suggesting a potential aberrant expression of DNA repair pathways in cRAG leukemic cells (Supplementary Figure 3B). These findings highlight the potential of cRAG to foster increased hybrid joint formation, especially when a normal pathway for efficient coding and RSS joining is compromised in an NHEJ-aberrant context.

Our data demonstrates that the deletion of the RAG1 non-core region results in more severe off-target V(D)J recombination compared to the deletion of the RAG2 non-core region. This observation is supported by the fact that the RAG1 terminus contains multiple zinc-binding motifs and ubiquitin ligase activity, which are known to enhance the efficiency of the rearrangement reaction (***Beilinson, et al., 2021;Burn, et al., 2022)***. Furthermore, our research reveals that RAG1 expression persists in *BCR-ABL1*^+^ progenitor B cells, and deletion of the non-core region of RAG1 results in elevated expression of RAG in comparison to fRAG. Consequently, as demonstrated in this study, cRAG1 from *BCR-ABL1*^+^ B leukemic mice is prone to generating off-target V(D)J recombination. The distinct function of RAG1’s non-core region in thymic lymphomas of Tp53-/- mice and *BCR-ABL1*^+^ B leukemic mice leads to dissimilar off-target activity of cRAG1 (***Mijuškovi***ć***, et al., 2015)***. Therefore, it would be intriguing to replicate these analyses across various subtypes of ALL to further investigate this phenomenon.

In human *ETV6-RUNX1* ALL, the *ETV6-RUNX1* fusion gene is believed to initiate prenatally, yet the disease remains clinically latent until critical secondary events occur, leading to leukemic transformation-“pre-leukemia to leukemia” (***Mijuškovi***ć***, et al., 2015)***. Genomic rearrangement, mediated by aberrant RAG recombinase activity, is a frequent driver of these secondary events in *ETV6-RUNX1* ALL (***Chen, et al., 2021;Papaemmanuil, et al., 2014)***. In contrast, RAG mediated off-target V(D)J recombination is also observed in *BCR-ABL1*^+^ B-ALL. These oncogenic structural variations can also be considered as secondary events that promote the transition - “leukemia to aggressive leukemia”. The enhancement of *BCR-ABL1*^+^ B-ALL deterioration and progression by cRAG in mouse model was consistent with our previous study that RAG enhances BCR-ABL1 positive leukemic cell growth through its endonuclease activity (***Yuan, et al., 2021)***. Additionally, we showed that non-core RAG1 region deletion leads to increased cRAG1 expression and high RAG expression related to low survival in pediatric acute lymphoid leukemia (Figure 3A and Supplementary Figure 7). Therefore, more attention should be paid to the non-core RAG region mutation in *BCR-ABL1*^+^ B-ALL for the role of non-core region in leukemia suppression and off-target V(D)J recombination.

## Methods

### Mice

The C57BL/6 mice were purchased from the Experimental Animal Center of Xi’an Jiaotong University, while cRAG1 (amino acids 384-1040) and cRAG2 (amino acids 1-383) were obtained from Dr. David G. Schatz (Yale University, New Haven, Connecticut, USA). The mice were bred and maintained in a specific pathogen-free (SPF) environment at the Experimental Animal Center of Xi’an Jiaotong University. All animal-related procedures were in accordance with the guidelines approved by the Xi’an Jiaotong University Ethics Committee for Animal Experiments.

### Generation of Retrovirus Stocks

The pMSCV-BCR-BAL1-IRES-GFP vector is capable of co-expressing the human BCR-ABL1 fusion protein and green fluorescence protein (GFP), while the pMSCV-GFP vector serves as a negative control by solely expressing GFP. To produce viral particles, 293T cells were transfected with either the MSCV-BCR-BAL1-IRES-GFP or MSCV-GFP vector, along with the packaging vector PKAT2, utilizing the X-tremeGENE HP DNA Transfection Reagent from Roche (Basel, Switzerland). After 48 hours, the viral supernatants were collected, filtered, and stored at −80°C.

### Bone Marrow Transduction and Transplantation

Experiments were conducted using mice aged between 6 to 10 weeks. *BCR-ABL1*^+^ B-ALL murine model was induced by utilizing marrow from donor mice who had not undergone 5-FU treatment. The donor mice were euthanized through CO_2_ asphyxiation, and the bone marrow was harvested by flushing the femur and tibia with a syringe and 26-gauge needle. Erythrocytes were not removed, and 1 × 10^6^ cells per well were plated in six-well plates. A single round of co-sedimentation with retroviral stock was performed in medium containing 5% WEHI-3B-conditioned medium and 10 ng/mL IL-7 (Peprotech, USA). After transduction, cells were either transplanted into syngeneic female recipient mice (1 × 10^6^ cells each) that had been lethally irradiated (2 × 450 cGy), or cultured in RPMI-1640 (Hyclone, Logan, UT) medium supplemented with 10% fetal calf serum (Hyclone), 200 mmol/L L-glutamine, 50 mmol/L 2-mercaptoethanol (Sigma, St Louis, MO), and 1.0 mg/mL penicillin/streptomycin (Hyclone). Subsequently, recipient mice were monitored daily for indications of morbidity, weight loss, failure to thrive, and splenomegaly. Weekly assessment of peripheral blood GFP percentage was done using FACS analysis of tail vein blood. Hematopoietic tissues and cells were utilized for histopathology, in vitro culture, FACS analysis, secondary transplantation, genomic DNA preparation, protein lysate preparation, or lineage analysis, contingent upon the unique characteristics of mice under study.

### Secondary Transplants

Thawed BM cells were sorted using a BD FACS Aria II (Becton Dickinson, San Jose California, USA). GFP positive leukemic cells (1× 10^6^, 1×10^5^, 1×10^4^, and 1×10^3^) were then resuspended in 0.4 mL Hank’s Balanced Salt Solution (HBSS) and intravenously administered to unirradiated syngeneic mice.

### Flow cytometry analysis and sorting

Bone marrow, spleen cells, and peripheral blood were harvested from leukemic mice. Red blood cells were eliminated using NH4Cl RBC lysis buffer, and the remaining nucleated cells were washed with cold PBS. In order to conduct in vitro cell surface receptor staining, 1× 10^6^ cells were subjected to antibody staining for 20 minutes at 4°C in 1×PBS containing 3% BSA. Cells were then washed with 1×PBS and analyzed using a CytoFLEX Flow Cytometer (Beckman Coulter, Miami, FL) or sorted on a BD FACS Aria II. Apoptosis was analysed by resuspending the cells in Binding Buffer (BD Biosciences, Baltimore, MD, USA), and subsequent labeling with antiannexin V-AF647 antibody (BD Biosciences) and propidium iodide (BD Biosciences) for 15 minutes at room temperature. The lineage analysis was performed using the following antibodies, which were purchased from BD Biosciences: anti-BP-1-PerCP-Cy7, anti-CD19-PerCP-CyTM^5.5^, anti-CD43-PE, anti-B220-APC, and anti-μHC-APC.

### BrdU incorporation and analysis

Cells obtained from primary leukemic mice were cultured in six-well plates containing RPMI-1640 medium supplemented with 10% FBS and 50 µg/mL BrdU. After incubation at 37°C for 30 minutes, the cells were harvested and intranuclearly stained with anti-BrdU and 7-AAD antibodies, following the manufacturer’s instructions.

### The in vitro V(D)J recombination assay

The retroviral recombination substrate pINV-12/23 was introduced into primary leukemic cells utilizing X-treme GENE HP DNA Transfection Reagent (Roche). Recombina-tion efficiency of pINV-12/23 was evaluated through flow cytometry analysis for mouse CD90 (mCD90) and hCD4 expression (***Yuan, et al., 2021)***.

### Western blotting analysis

Over 1 × 10^6^ leukemic cells were centrifuged and washed with ice-cold PBS. The cells were then treated with ice-cold RIPA buffer, consisting of 50 mM Tris-HCl (pH 7.4), 0.15 M NaCl, 1% Triton X-100, 0.5% NaDoc, 0.1% sodium dodecyl sulphide (SDS), 1 mM ethylene diamine tetraacetic acid (EDTA), 1 mM phenylmethane sulphony fluoride (PMSF) (Amresco), and fresh protease inhibitor cocktail Pepstain A (Sigma). After sonication using a Bioruptor TMUCD-200 (Diagenode, Seraing, Belgium), the suspension was spined at 14,000 g for 3 minutes at 4°C. The total cell lysate was either utilized immediately or stored at −80°C. Protein concentrations were determined using DC Protein Assay (Bio–Rad Laboratories, Hercules, California, USA). Subsequently, the protein samples (20 μg) were incubated with α-RAG1 (mAb 23) and α-RAG2 (mAb 39) antibodies (***Teng, et al., 2015)***, with GAPDH serving as the loading control. The signal was further detected using secondary antibody of goat anti-rabbit IgG conjugated with horseradish peroxidase (Thermo Scientific, Waltham, MA). The band signal was developed with Immobilon™ Western Chemiluminescent HRP substrate (Millipore, Billerica, MA). The band development was analyzed using GEL-PRO ANALYZER software (Media Cybernetics, Bethesda, MD).

### Genomic PCR

Genomic PCR was performed in a 20μl reaction containing 50 ng of genomic DNA, 0.2 μm of forward and reverse primer, and 10 μl Premix Ex Taq (TaKaRa, Shiga, Japan). Amplification conditions were as follows: 94°C for 5 minutes; 35 cycles of 30 seconds at 94°C, 30 seconds at 60°C and 1 minutes at 72°C; 72°C for 5 minutes (BioRad, Hercules, CA). Genomic PCR was performed using the following primers: D_H_L-5′-GGAATTCGMTTTTTGTSAAGGGATCTACTACTGTG-3′;J_H_3-5′-GTCTAGATTCTCAC-AAGAGTCCGATAGACCCTGG-3′; V_H_Q52-5′-CGGTACCAGACTGARCATCASCAAG-GACAAYTCC-3′; V_H_558-5′-CGAGCTCTCCARCACAGCCTWCATGCARCTCARC-3′; V_H_7183-5′-CGGTACCAAGAASAMCCTGTWCCTGCAAATGASC-3′.(***Schlissel, et al., 1991)***:

### RNA-seq library preparation and sequencing

GFP^+^CD19^+^ cells were sorted from the spleen of cRAG1 (n=3, 1×10^6^ cells /sample), cRAG2 (n=3, 1×10^6^ cells /sample), and fRAG (n=3, 1×10^6^ cells /sample) *BCR-ABL1*^+^ B-ALL mice. Total RNA was extracted using Trizol reagent (Invitrogen, CA, USA) following the manufacturer’s guidelines. RNA quantity and purity analysis was done using Bioanalyzer 2100 and RNA 6000 Nano LabChip Kit (Agilent, CA, USA) with RIN number >7.0. RNA-seq libraries were prepared by using 200 ng total RNA with TruSeq RNA sample prep kit (Illumina). Oligo(dT)-enriched mRNAs were fragmented randomly with fragmentation buffer, followed by first- and second-strand cDNA synthesis. After a series of terminal repair, the double-stranded cDNA library was obtained through PCR enrichment and size selection. cDNA libraries were sequenced with the Illumina Hiseq 2000 sequencer (Illumina HiSeq 2000 v4 Single-Read 50 bp) after pooling according to its expected data volume and effective concentration. Two biological replicates were performed in the RNA-seq analysis. Raw reads were then aligned to the mouse genome (GRCm38) using Tophat2 RNA-seq alignment software, and unique reads were retained to quantify gene expression counts from Tophat2 alignment files. The differentially expressed mRNAs and genes were selected with log2 (fold change) >1 or log2 (fold change) <-1 and with statistical significance (p value < 0.05) by R package. Bioinformatic analysis was performed using the OmicStudio tools athttps://www.omicstudio.cn/tool.

### Preparation of tumor DNA samples

GFP^+^CD19^+^ splenic cells, tail and kidney tissue were obtained from *cRAG1, cRAG2* and *fRAG BCR-ABL1*^+^ B-ALL mice, and genomic DNA was extracted using a TIANamp Genomic DNA Kit (TIANGEN-DP304). Subsequently, paired-end libraries were constructed from 1 µg of the initial genomic material using the TruSeq DNA v2 Sample Prep Kit (Illumina, #FC-121-2001) as per the manufacturer’s instructions. The size distribution of the libraries was assessed using an Agilent 2100 Bioanalyzer (Agilent Technologies, #5067-4626), and the DNA concentration was quantified using a Qubit dsDNA HS Assay Kit (Life Technologies, #Q32851). The Illumina HiSeq 4000 was utilized to sequence the samples, with two to four lanes allocated for sequencing the tumor and one lane for the control DNA library of the kidney or liver, each with 150 bp paired end reads.

### Read alignment and structural variant calling

Fastq files were generated using Casava 1.8 (Illumina), and BWA 37 was employed to align the reads to mm9. PCR duplicates were eliminated using Picard’s Mark Duplicates tool *(source-forge.net/apps/mediawiki/picard)*. Our custom scripts (http://sourceforge.net/projects/svdetection) were utilized to eliminate BWAdesignated concordant and read pairs with low BWA mapping quality scores. Intrachromosomal and inter-chromosomal rearrangements were identified using SV Detect from discordant, quality prefiltered read pairs. The mean insertion size and standard deviation for this analysis were obtained through Picard’s InsertSizeMetrics tool (sourceforge.net/apps/mediawiki/picard). Tumor-specific structural variants (SVs) were identified using the manta software *(https://github.com/Illumina/manta/blob/mater/docs/userGuide/README.md#introduction)*.

### Validation of high confidence off-target candidates

The elimination of non-specific structural mutations from the kidney or tail was necessary for tumor-specific structural variants identification. Subsequently, the method involving 21-bp CAC-to-breakpoint was employed to filter RAG-mediated off-target gene. The validation of high confidence off-target candidates was carried out through PCR. Oligonucleotide primers were designed to hybridize within the “linking” regions of SV Detect, in the appropriate orientation. The PCR product was subjected to Sanger sequencing and aligned to the mouse mm9 reference genome using BLAST (https://blast.ncbi.nlm.nih.gov/Blast.cgi).

### Statistics

Statistical analysis was conducted using SPSS 20.0 (IBM Corp.) and GraphPad Prism 6.0 (GraphPad Software). Descriptive statistics were reported as means ± standard deviation for continuous variables. Statistical analyses were applied to biologically independent mice or technical replicates for each experiment which was independently repeated at least three times. The equality of variances was assessed using Levene’s test. Two-group comparisons, multiple group comparisons, and survival comparisons were performed using independent-samples t-test, one-way analyses of variance (ANOVA) with post hoc Fisher’s LSD test, and log-rank Mantel-Cox analysis, respectively. Kaplan-Meier survival curves were utilized to depict the changes in survival rate over time. Statistical significance was set at *P*<0.05.

## Supporting information

Revise figure legends

## Authors’ Contributions

Yanghong Ji: Conceptualization, resources, data curation, funding acquisition, validation, writing-review, and editing. Xiaozhuo Yu and Wen Zhou: Conceptualization, validation, visualization, methodology, writing-original draft, writing-review, and editing. Xiaodong Chen: validation, writing-review, and editing. Shunyu He: methodology, writing-review, and editing. Mengting Qin: writing-review, and editing. Meng Yuan: validation, writing-review, and editing. Yang Wang: validation, writing-review and editing. Woodvine otieno Odhiambo: writing-review and editing. YinSha Miao: funding, validation, writing-review, and editing.

## Acknowledgments

This study was supported by grants (no. 31170821, no. 31370874 and no. 81670157) from the National Natural Scientific Foundation of China and by a grant (no. 2016JZ0-30) from the Natural Scientific Foundation of Shaanxi. The authors would like to thank Professor Shaoguang Li from the Division of Hematology/Oncology, University of Massachusetts Medical School, for providing the MSCV-BCR-BAL1-IRES-GFP construct. The authors would also like to thank Mr. Xiaofei Wang (Xi’an Jiaotong University Health Science Centre) for providing expert technical assistance with cell sorting.

