## Supplementary figures and images for "RAG1 and RAG2 Non-core Regions Are Implicated in Leukemogenesis and Off-target V(D)J Recombination in BCR-ABL1-driven B-cell Lineage Lym-phoblastic Leukemia"

### supplementary figure 1

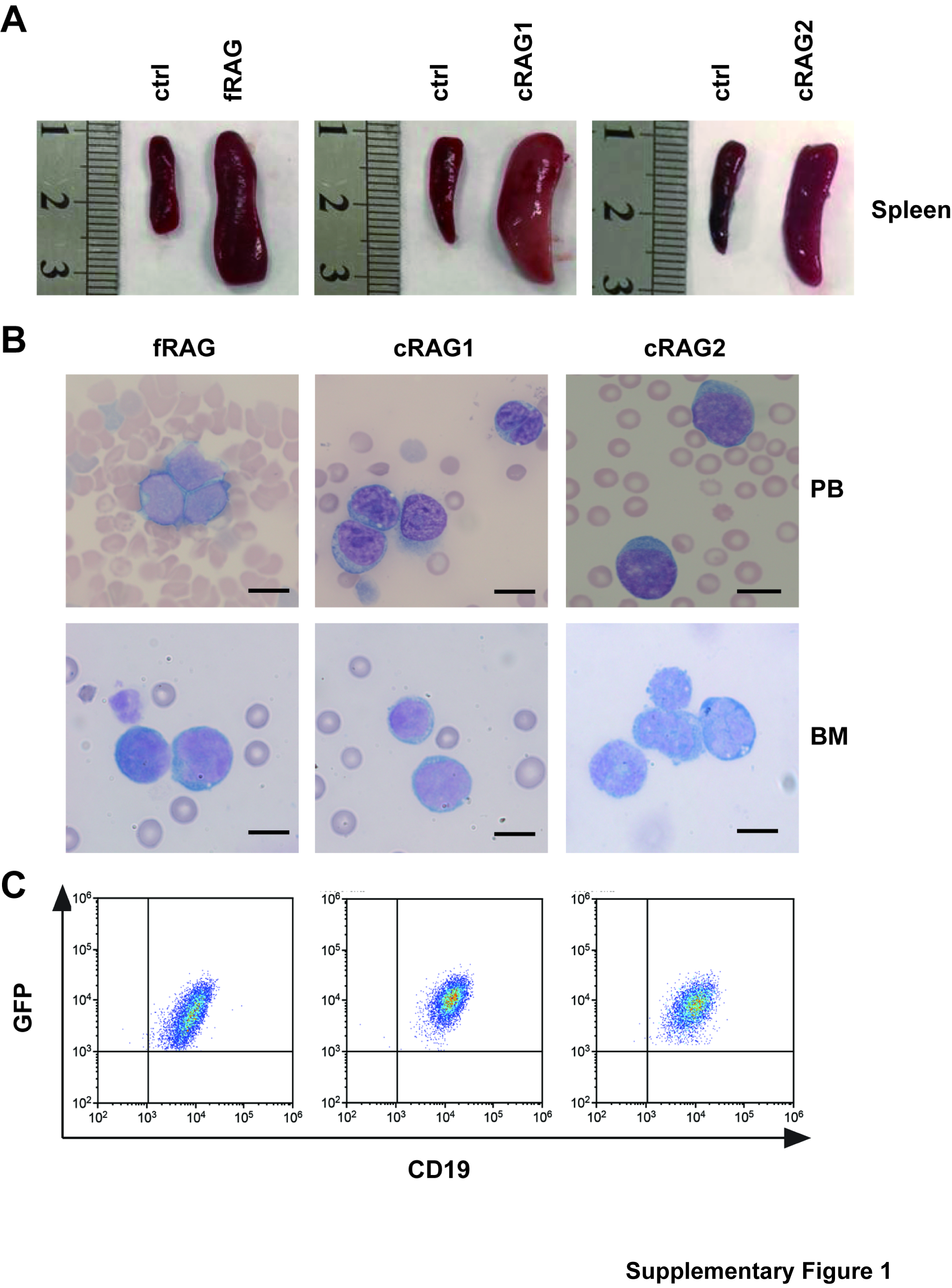

### supplementary figure 2

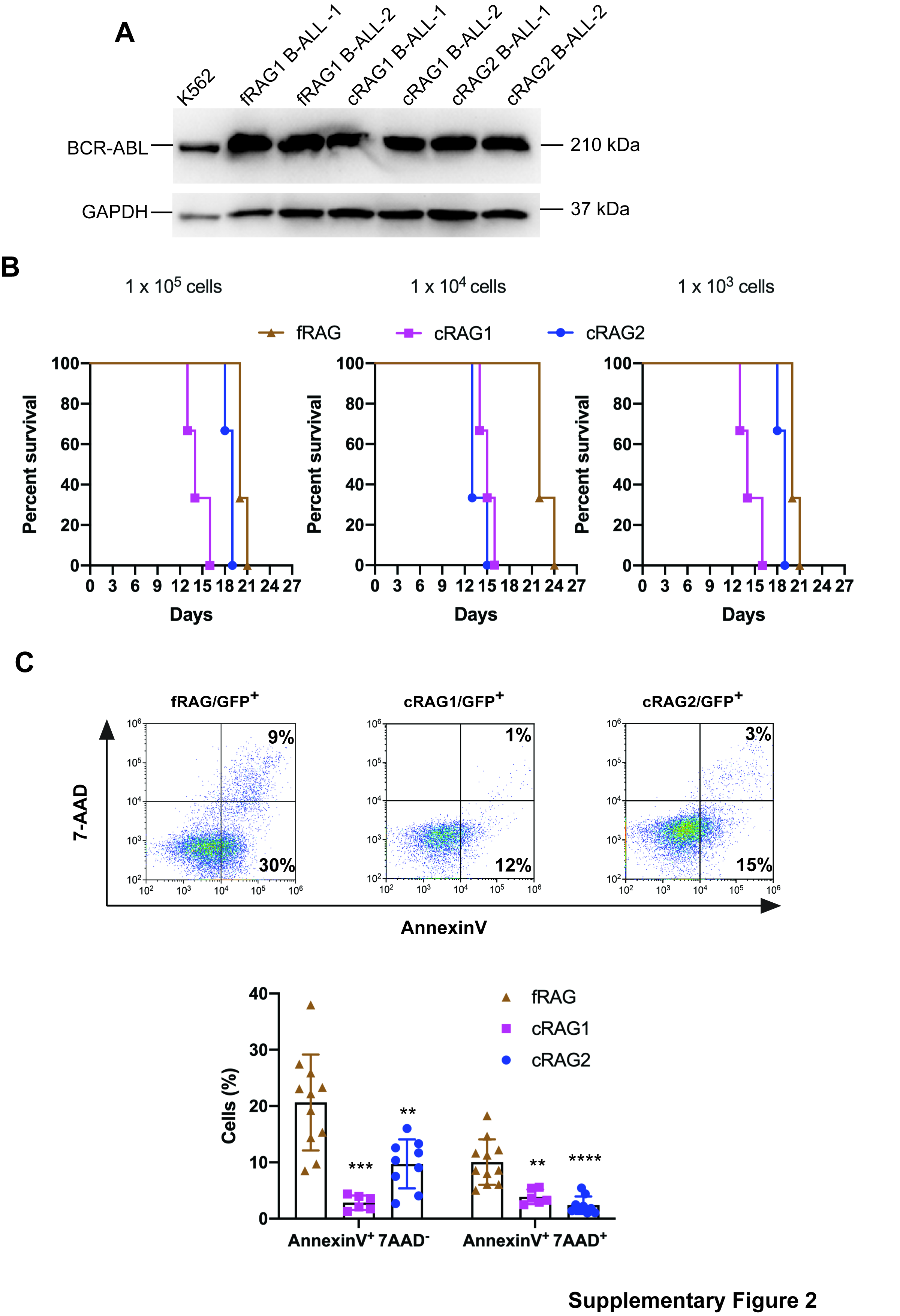

### supplementary figure 3

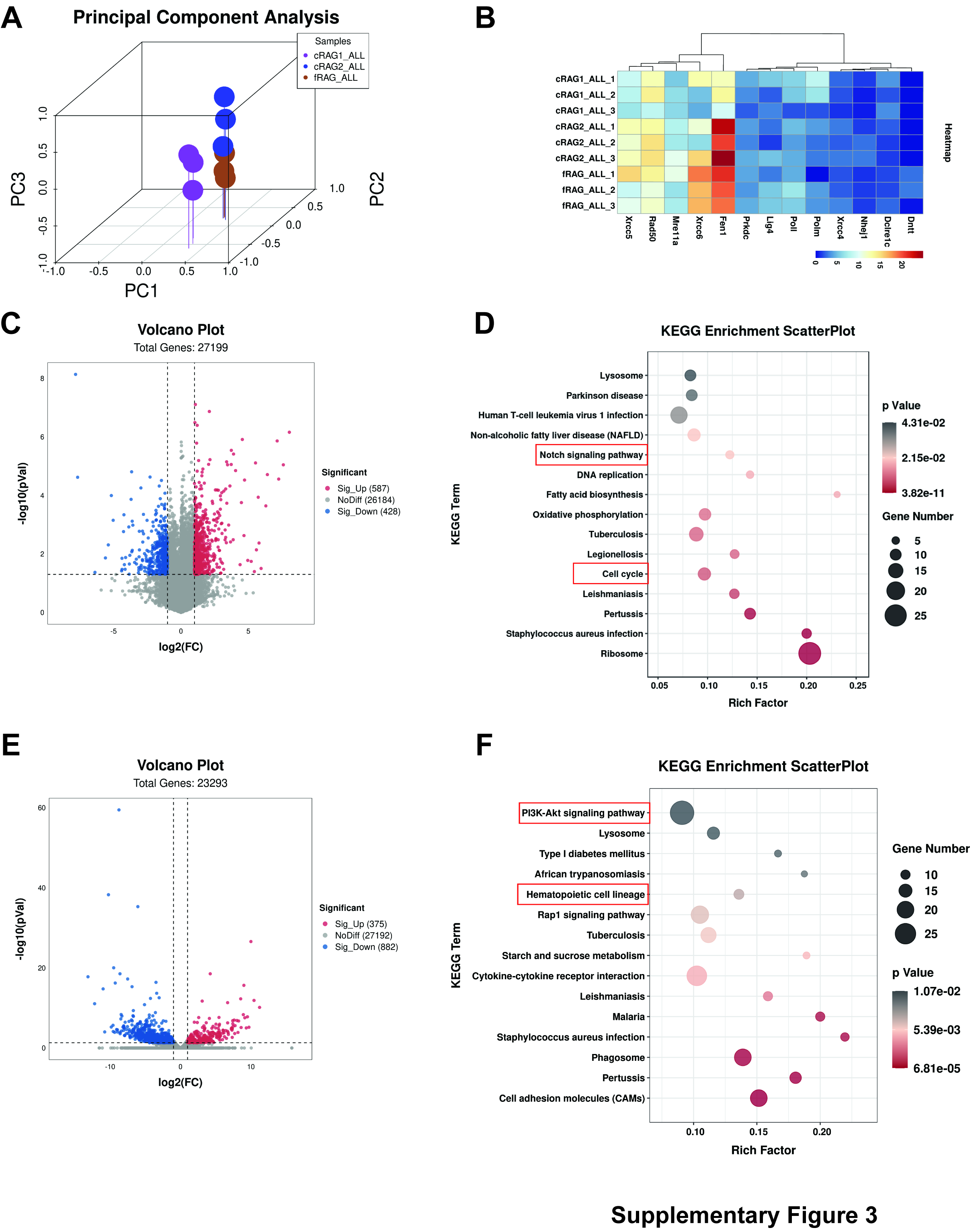

### supplementary figure 4

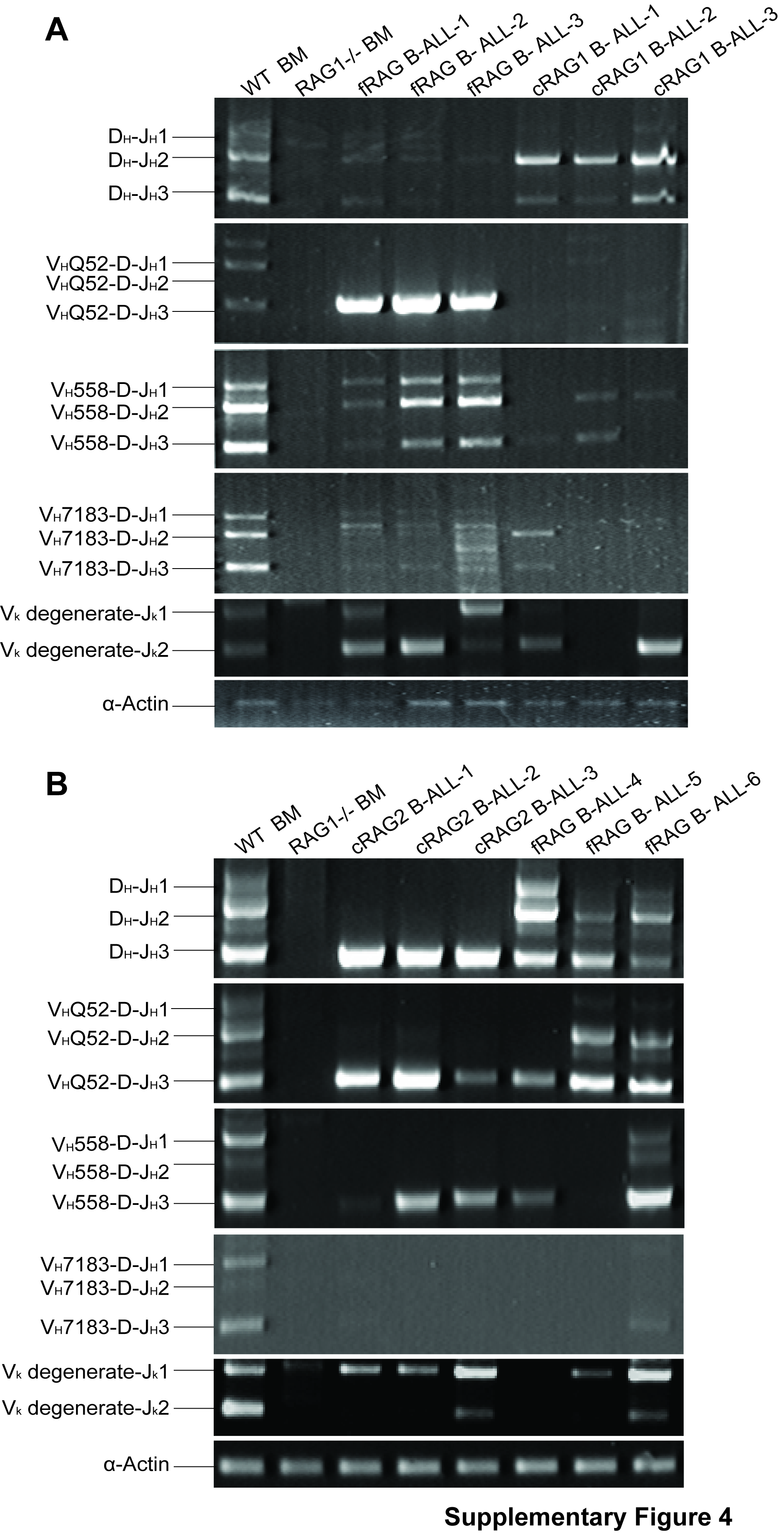

### supplementary figure 5

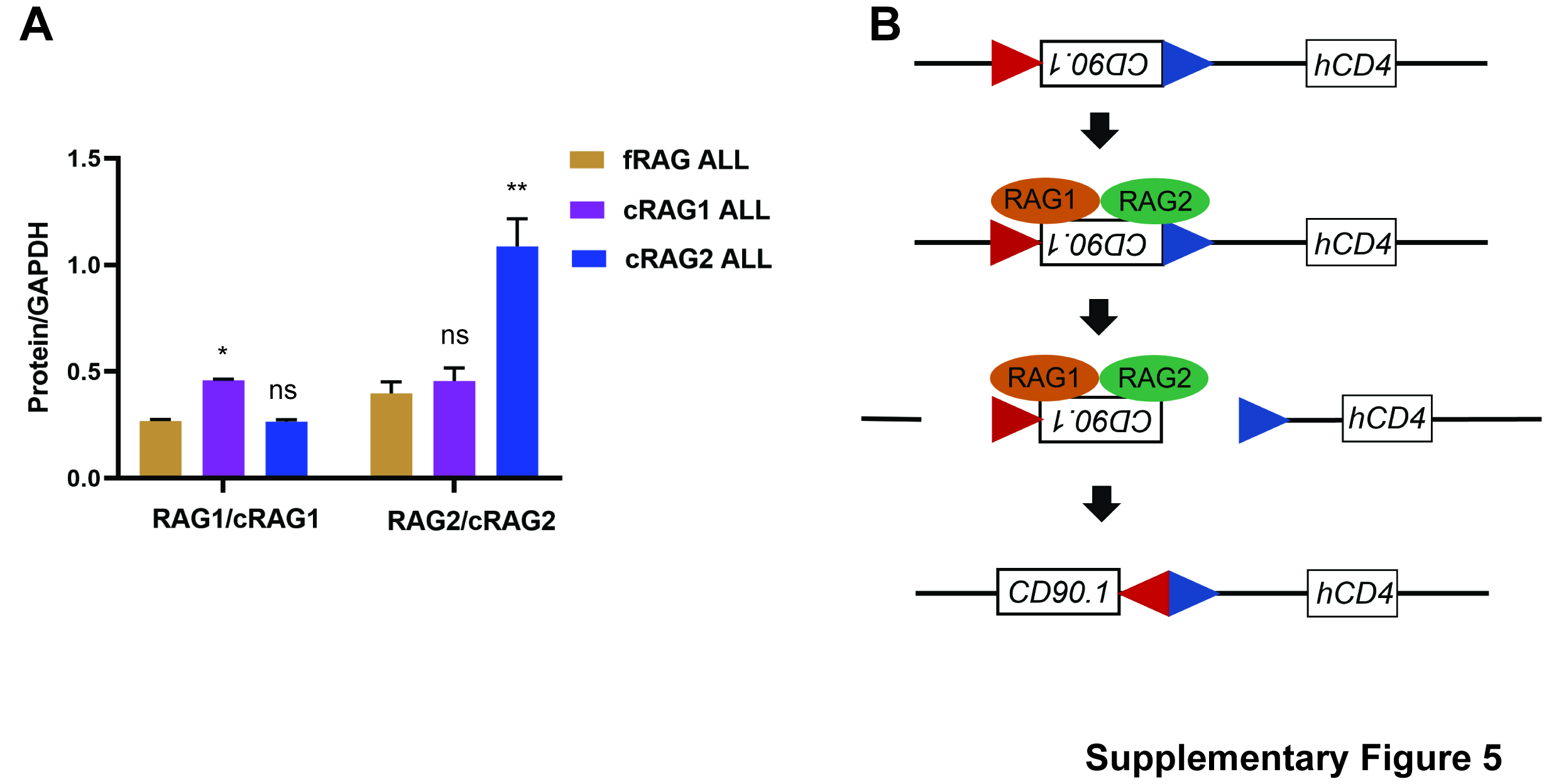

### supplementary figure 6

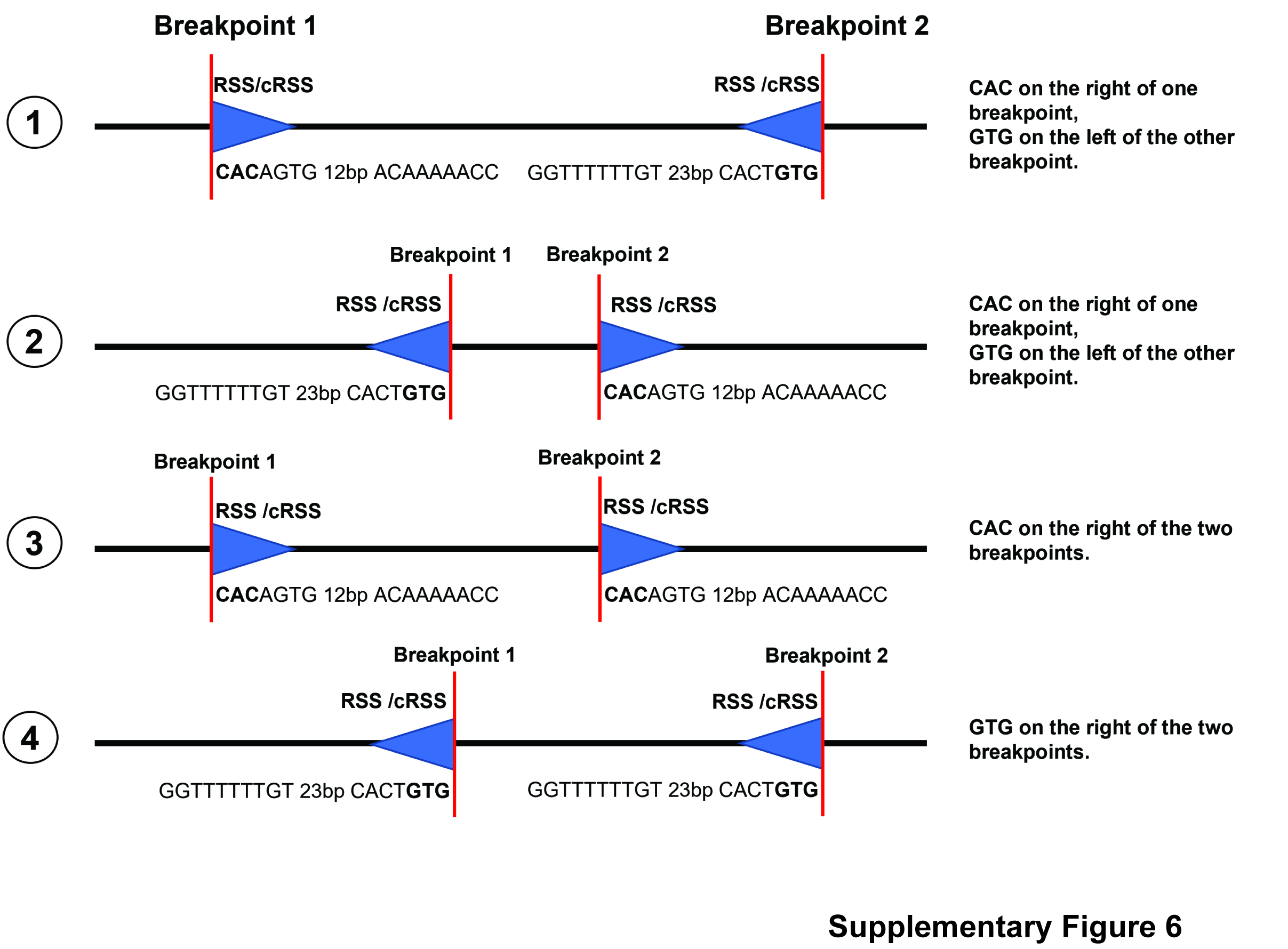

### supplementary figure 7

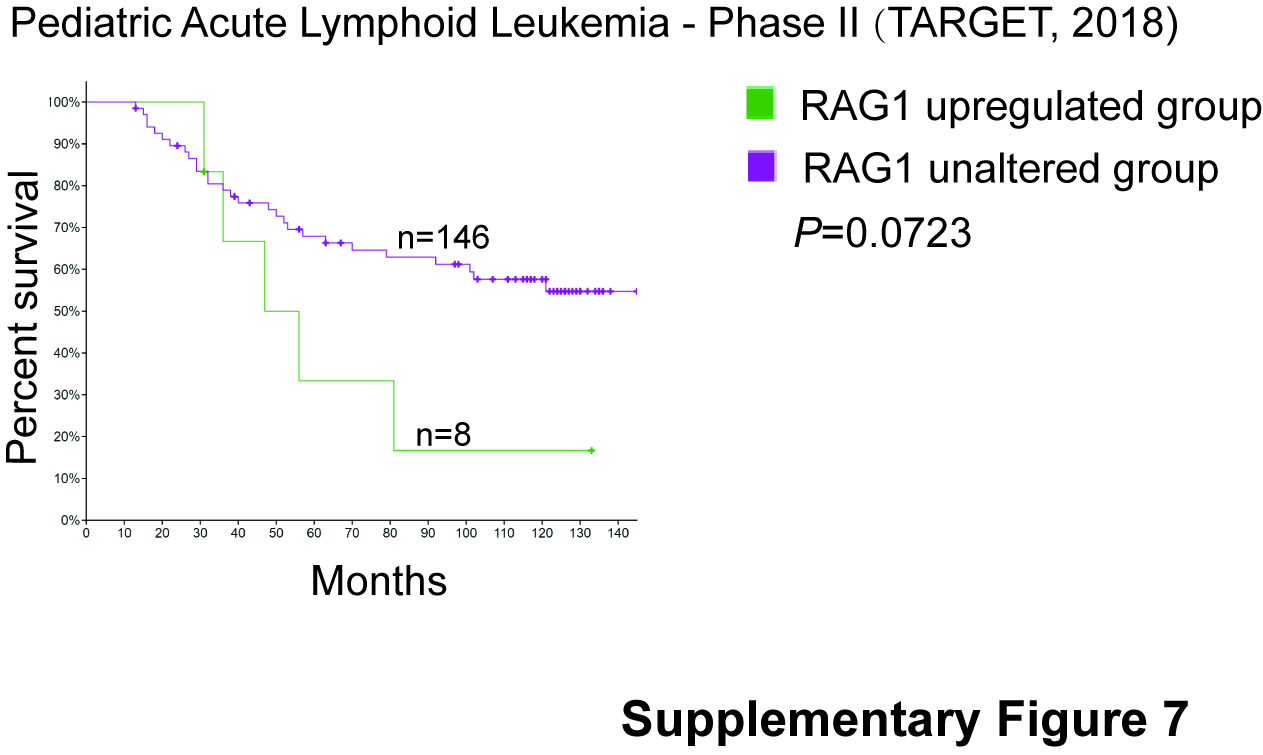
