## Supplementary material for "RAG1 and RAG2 Non-core Regions Are Implicated in Leukemogenesis and Off-target V(D)J Recombination in BCR-ABL1-driven B-cell Lineage Lym-phoblastic Leukemia": Revise figure legends

**Figure legend**

**Figure 1. cRAGs give more aggressive leukemia in mice model of *BCR-ABL1*^+^ B-ALL**

(A) Kaplan-Meier survival curve for fRAG (n=8), cRAG1 (n=6), and cRAG2 (n=10) recipient mice. The survival was calculated by Mantel–Cox test (fRAG vs cRAG1, P<0.0381, fRAG vs cRAG2, *P*<0.0412). (B) The spleen weights of fRAG*,* cRAG1 and cRAG2 leukemic mice (fRAG, n=8, cRAG1, n=7, cRAG2, n=9; fRAG vs cRAG1, *P*<0.0001, fRAG vs cRAG2, *P*=0.1352). (C) The numbers of spleen cell in fRAG*,* cRAG1 and cRAG2 leukemic mice (fRAG, n=7, cRAG1, n=8, cRAG2, n=13; fRAG vs cRAG1, *P*=0.0047, fRAG vs cRAG2, *P*=0.0180). (D) The percentage of GFP^+^ cells in peripheral blood (PB), (fRAG, n=6, cRAG1, n=6, cRAG2, n =6; fRAG vs cRAG1, *P*=0.0003, fRAG vs cRAG2, *P*=0.0035), bone marrow (BM, fRAG, n=5, cRAG1, n=5, cRAG2, n=6; fRAG vs cRAG*1*, *P*=0.0341, fRAG vs cRAG2, *P*=0.0008), and spleen (SP, fRAG, n=9, cRAG1, n=4, cRAG2, n=9; fRAG vs cRAG1, *P*=0.0016, fRAG vs cRAG2, *P*<0.0001) of fRAG*,* cRAG1 and cRAG2 leukemic mice. (E) Representative flow cytometry plots of cell cycle arrest of leukemic cells in fRAG*,* cRAG1 and cRAG2 mice. In the graph, the percentages of each phase of the cell cycle are summarized below (fRAG, n=3, cRAG1, n=5, cRAG2, n=5; G0/G1, fRAG vs cRAG1, *P*=0.0082, fRAG vs cRAG2, *P*=0.0279; S, fRAG vs cRAG1, *P*=0.0146, fRAG vs cRAG2, *P*=0.0370; G2/M, fRAG vs cRAG1, *P*=0.0134, fRAG vs *cRAG2*, *P*=0.1507). In figures B, C, D and J, error bars represent the mean ± s.d., *P* values were calculated by Student’s t test and **P* < 0.05, ***P* < 0.01, ****P* < 0.001.

**Figure 2. The non-core RAG region loss corresponds to a less mature cell surface phenotype**

(A) Flow cytometry analysis of the B cell markers CD19, BP-1, B220, and CD43 on *BCR-ABL1*-transformed fRAG, cRAG1 and cRAG2 leukemic bone marrow cells. The percentages of each phase of the B cell stage are summarized in the bottom graph (fRAG, n=9, cRAG1, n=4, cRAG2, n=9; Large-preB, fRAG vs cRAG1, *P*=0.0349, fRAG vs cRAG2, *P*=0.0017; Small-pre-B, fRAG vs cRAG1, *P*=0.0141, fRAG vs cRAG2, *P*=0.0005). The expression of the cytoplasmic μ chain was analyzed by flow cytometry. Representative samples are shown in (B), and the results from multiple samples analyzed in independent experiments are summarized in the bottom graph as the fraction of cells expressing cytoplasmic factors (fRAG, n=11, cRAG1, n=9, cRAG2, n=8; fRAG vs cRAG1, *P*=0.0312, fRAG vs cRAG2, *P*=0.0441). Error bars represent the mean ± s.d., *P* values were calculated by Student’s t test and **P* < 0.05, ***P* < 0.01, ****P* < 0.001.

**Figure 3. The non-core RAG region loss highlights genomic DNA damage**

(A) Western blotting analysis showed RAG1 and RAG2 expression in GFP^+^CD19^+^ leukemic cells originating from *BCR-ABL1^+^* B-ALL in different genetic backgrounds. The experiment was repeated under the same conditions three times. (B) Rearrangement substrate retrovirus was transduced into leukemic cells. Flow cytometry was used to analyze the percentage of CD90.1 and hCD4 positive cells, and the percentage populations are shown in the bottom graph (fRAG, n=3, cRAG1, n=3, cRAG2, n=3; fRAG vs cRAG1, *P*=0.0002, fRAG vs cRAG2, *P*=0.5865). (C) Flow cytometry analysis of ɣ-H2AX levels in fRAG*,* cRAG1 and cRAG2 leukemic cells and the percentage of ɣ-H2AX-positive cell populations shown in the bottom graph (fRAG, n=11, cRAG1, n=8, cRAG2, n=8; fRAG vs cRAG1, *P*=0.0505, fRAG vs cRAG2, *P*=0.0094). Error bars represent the mean ± s.d., *P* values were calculated by Student’s t test and **P* < 0.05, ***P* < 0.01, ****P* < 0.001.

**Figure 4. Structural alterations in *BCR-ABL1*+ B lymphocytes**

(A-C) Circos plot representation of all off-target recombination detected in the genome-wide analyses of fRAG, cRAG1 and cRAG2 leukemic cells. See also Table S3.

**Figure 5. Overview and characteristics of off-target recombination in** ***BCR-ABL1^+^* B-ALL leukemic cells from *fRAG* and *cRAG* mice**

(A)Exon-intron distribution profiles of 41 breakpoints generated by 24 SVs. Gene body includes exon (n = 9; 17.3%) and intron (n = 20; 38.5%). Flanking sequence includes 3'UTR (n = 6; 11.5%), 5'UTR (n = 2; 3.8%), promoter (n = 6; 11.5%), and downstream (n = 9; 17.3%). (B) the off-target recombination was filtered and verified by whole genomic sequence and PCR respectively. P nucleotides and N nucleotides of RSS to RSS and cRSS to cRSS were calculated in *BCR-ABL1*^+^ B-ALL. (C) Hybrid joint percentage generated by either fRAG, cRAG1 or cRAG2 in *BCR-ABL1*^+^ B-ALL. It was 0, 100%, and 93% in fRAG, cRAG1 or cRAG2 leukemic cells respectively. (D)The 24 off-target recombination genes were retrieved by COSMIC Cancer Gene Census *(*[*http://cancer.sanger.ac.uk/census/*](http://cancer.sanger.ac.uk/census/)*)*. 0.5 genes and 0.5 cancers gene average sample in fRAG leukemic cells; 8 genes and 4.5 cancer genes average sample in cRAG1 leukemic cells; 3.3 genes and 0.3 cancer genes average sample in cRAG2 leukemic cells.

### Figure 6. The non-core regions have effects on RAG binding accuracy and recombinat size in *BCR-ABL1^+^* B lymphocyte

(A) Sequence logos were used to compare the RSS and cRSS in Ig loci and non-Ig loci. Top panel: V(D)J recombination at *Ig* locus; the next three panels: RAG-mediated off-target recombination at non-Ig locus from fRAG, cRAG1 and cRAG2 leukemic cells respectively. The scale of recombinant size was categorized into three ranges: <1000bp, 1000-10000bp, and >10000bp. The distribution of different recombinant sizes in fRAG, cRAG1, and cRAG2 leukemic cells was presented in (B), while the number of different recombinant sizes in fRAG, cRAG1, and cRAG2 leukemic cells was displayed in (C). (D) A schematic depiction of the mechanism of cRAG-accelerated off-target V(D)J recombination was provided. Both RAG1 and RAG2’s non-core region deletion decreases RAG binding accuracy in cRAG1 and cRAG2, *BCR-ABL1*^+^ B ALL. Additionally, RAG1’s non-core region deletion significantly reduces the size and scale of off-target V(D)J recombination in cRAG1, *BCR-ABL1*^+^ B ALL.

### Supplementary Figure 1. Construction of fRAG, cRAG1 and cRAG2*,* *BCR-ABL1^+^* B-ALL mice models using bone marrow transplantation (BMT)

In the establishment of *BCR-ABL1*^+^ B-ALL mice models, fRAG, cRAG1 and cRAG2 recipient mice after syngeneic lethal irradiation were transplanted with corresponding donor bone marrow cells transduced by *MSCV-BCR-BAL1-IRES-GFP* or *MSCV-GFP* retroviral supernatants. (A) Gross appearance of the spleen in fRAG, cRAG1, and cRAG2 leukemic mice and corresponding control mice. (B) Peripheral blood (PB) and bone marrow (BM) lymphoblastic cells were stained by Wright-Giemsa. The scale bars represent 10 µm. (C) Bone marrow cells from fRAG, cRAG1 and cRAG2 leukemic mice were examined by flow cytometry for the expression of GFP and CD19.

**Supplementary Figure 2. Biological behavior of leukemia in fRAG, cRAG1 and cRAG2 BCR-ABL1+ B-ALL mouse model**

(A) BCR-ABL1 expressions in GFP^+^CD19^+^ leukemic cells were determined by western. GAPDH protein was used as a loading control. The K562 and 293T cell lines served as the positive control and negative controls, respectively. (B) Survival of secondary transplant setting. Leukemia cells from primary recipients were recovered from the spleens and purified by GFP^+^ cell sorting. A total of 10^5^,10^4^ and 10^3^ GFP^+^ leukemia cells originating from fRAG, cRAG1 or cRAG2 *BCR-ABL1^+^* B-ALL were transplanted into corresponding nonirradiated immunocompetent syngenetic recipient mice (fRAG, n=3, cRAG1, n=3, cRAG2, n=3; survival days fRAG, 11-26 days, cRAG1, 10-16 days, cRAG2, 11-21 days in three different concentrations of GFP^+^ leukemic cells; fRAG vs cRAG1, *P*=0.0299, fRAG vs cRAG2, *P*=0.2286 in concentrations of 10^5^, fRAG vs cRAG1, *P*=0.0246, fRAG vs cRAG2, *P*=0.0295 in concentrations of 10^4^, fRAG vs cRAG1, *P*=0.0246, fRAG vs cRAG2, *P*=0.0224 in concentrations of 10^3^, by Mantel–Cox test). (C) Apoptosis was measured by flow cytometry (Annexin V and 7-AAD). The Annexin V^+^ and 7-AAD^-^ cells were defined as early apoptotic cells, while Annexin V^+^ and 7-AAD^+^ cells were late apoptotic cells (fRAG, n=11, cRAG1, n=6, cRAG2, n=9; early apoptotic cells: fRAG vs cRAG1, *P*=0.0002, fRAG vs cRAG2, *P*=0.0026; late apoptotic cells, fRAG vs cRAG1, *P*=0.0026, fRAG vs cRAG2, *P*<0.0001). Error bars represent the mean ± s.d. *P* values were calculated by Student’s t test and **P* < 0.05, ***P* < 0.01, ****P* < 0.001, *****P* < 0.0001.

**Supplementary Figure 3. The genetic pathways in fRAG, cRAG1, and cRAG2 *BCR-ABL1*+ lymphocytes.**

mRNA sequence was performed in GFP and CD19 double positive cells. (A) Principal Component Analysis (PCA) showing the distribution of differentially expressed samples of fRAG, cRAG1, and cRAG2, *BCR-ABL1*^+^ B-ALL. (B) Heatmap of representative different expressed genes related to non-homologous end repair. The scale ranges from minimum (blue) to medium (yellow) to maximum (red) relative expression. (C) Volcano plot depicting log2 (fold change) (x-axis) and −log10 (*P* value) (y-axis) for differentially expressed genes (FC > 2, p < 0.05) in GFP^+^CD19^+^ leukemic cells sorted from fRAG and cRAG1*, BCR-ABL1*^+^ B-ALL mice; upregulated (red) and downregulated (blue). n = 3 per group. (D) The Kyoto Encyclopedia of Genes and Genomes (KEGG) analysis was conducted to identify the differentially expressed genes in cRAG1 *BCR-ABL1*^+^ B-ALL. The top 15 pathways that exhibited significant differences were listed in this paragraph. The cell proliferation, apoptosis, and differentiation related pathway were highlighted in red squares. (E) Volcano plot depicting log2 (fold change) (x-axis) and −log10 (*P* value) (y-axis) for differentially expressed genes (FC > 2, *P* < 0.05) in GFP^+^CD19^+^ leukemic cells sorted from fRAG and cRAG2*, BCR-ABL1^+^* B-ALL mice; upregulated (red) and downregulated (blue). n = 3 per group. (D) KEGG analysis was conducted to identify the differentially expressed genes in cRAG2 *BCR-ABL1*^+^ ALL. The top 15 pathways that exhibited significant differences were listed in this paragraph. The cell proliferation, apoptosis, and differentiation related pathway were highlighted in red squares.

**Supplementary Figure 4. VDJ recombination in leukemic cells with different genetic backgrounds**

(A) VDJ recombination was analyzed by genomic PCR in GFP^+^CD19^+^ cell’s DNA from fRAG, cRAG1 or cRAG2 leukemic cells*.* Genomic DNA from RAG1-/- bone marrow cells and WT spleen was used as negative and positive control respectively.

**Supplementary Figure 5. RAG protein expression levels and schematic diagram of the recombinant substrate vector**

(A)RAG1/cRAG1 and RAG2/cRAG2 protein levels were compared by western blot and ImageJ software in fRAG1, cRAG1 and cRAG2, B-ALL cells. Error bars represent the mean ± s.d. The *P* value was calculated by t test, ***P*<0.01, **P*<0.05, ns *P*>0.05. (B) The B-ALL cells were subjected to transformation with the recombinant substrate vector. In the event of expression of RAG recombinase in the leukemic cells, the RSS sequences flanking CD90.1 would be cleaved by RAG, thereby facilitating the positioning and expression of both CD90.1 and hCD4. In the absence of RAG expression, only hCD4 would be expressed.

**Supplementary Figure 6. The criteria for identifying off-target recombination.**

We adopted the criteria used in previous studies. First, a CAC must exist to the right (or GTG to the left) of both breakpoints, which includes the four RAG-mediated DNA fragmentation cases mentioned above, and second, it must occur within a specified distance from the breakpoint and the CAC distance-to-breakpoint value was set at 21 bp.

**Supplementary Figure 7. The relationship of RAG1 mRNA levels and survival of pediatric acute lymphoid leukemia.**

The relationship of RAG1 mRNA levels and survival of pediatric acute lymphoid leukemia was research by cBioPortal *(https://www.cbioportal.org/)*. RAG1 mRNA levels were studied in pediatric patients with ALL at the time of diagnosis. The patients were separated into two groups based on mRNA levels of RAG1 (mRNA expression z score relative to diploid sample, RNA sequence RPKM, RAG1 upregulated group, n=8; RAG1 unaltered group, n=146). The *P* values were calculated from the log-rank test, *P*=0.0732.
